# Characterization of the skeletal muscle arginine methylome in health and disease reveals remodeling in Amyotrophic Lateral Sclerosis

**DOI:** 10.1101/2024.01.08.574551

**Authors:** Julian P. H. Wong, Ronnie Blazev, Yaan-Kit Ng, Craig A. Goodman, Magdalene K. Montgomery, Kevin I. Watt, Christian S. Carl, Matthew J. Watt, Christian T. Voldstedlund, Erik A. Richter, Peter J. Crouch, Frederik J Steyn, Shyuan T Ngo, Benjamin L. Parker

## Abstract

Arginine methylation is a protein post-translational modification important for the development of skeletal muscle mass and function. Despite this, our understanding of the regulation of arginine methylation under settings of health and disease remains largely undefined. Here, we investigated the regulation of arginine methylation in skeletal muscles in response to exercise and hypertrophic growth, and in diseases involving metabolic dysfunction and atrophy. We report a limited regulation of arginine methylation under physiological settings that promote muscle health, such as during growth and acute exercise, nor in disease models of insulin resistance. In contrast, we saw a significant remodeling of asymmetric dimethylation in models of atrophy characterized by the loss of innervation, including in muscle biopsies from patients with amyotrophic lateral sclerosis (ALS). Mass spectrometry-based quantification of the proteome and asymmetric arginine dimethylome of skeletal muscle from individuals with ALS revealed the largest compendium of protein changes with the identification of 793 regulated proteins, and novel site-specific changes in asymmetric dimethyl arginine (aDMA) of key sarcomeric and cytoskeletal proteins. Finally, we show that *in vivo* overexpression of PRMT1 and aDMA resulted in increased fatigue resistance and functional recovery in mice. Our study provides evidence for asymmetric dimethylation as a regulator of muscle pathophysiology and presents a valuable proteomics resource and rationale for numerous methylated and non-methylated proteins, including PRMT1, to be pursued for therapeutic development in ALS.

## INTRODUCTION

Skeletal muscles account for 30-40% of total body weight and are vital for the maintenance of posture, movement, and overall functional independence ^1^. In states of skeletal muscle disease (i.e. ALS or dystrophy) or disuse (i.e. immobilization or denervation), atrophy is common and reduces myofiber protein content, contractility and fatigue resistance, resulting in detrimental functional impairments in affected individuals ^2^. Skeletal muscles are also central regulators of whole-body metabolism by serving as primary sites for glucose uptake and storage. In turn, the desensitization of skeletal muscles to insulin (insulin resistance) is a major mechanism underpinning type-2 diabetes (T2D) ^3^. Maintaining skeletal muscle health through various exercise modalities is among the most effective treatments for muscle wasting and T2D ^4–8^, and further protects us from a myriad of other chronic conditions, such as cardiovascular diseases and premature death ^9,10^. Thus, a greater understanding of the signaling mechanisms that govern skeletal muscle in health and disease can enable the identification of novel therapeutic targets that could be pursued for the betterment of our well-being and longevity.

Over the last two decades, arginine methylation has emerged as a key protein post-translation modification (PTM) capable of modifying a diverse array of substrates ^11^. In mammals, protein arginine methylation is carried out by one of nine protein arginine methyltransferases (PRMT), resulting in the synthesis of three distinct forms of methylarginine: mono-methylarginine (MMA), asymmetric dimethyl arginine (aDMA) and symmetric dimethyl arginine (sDMA) ^11^. Previous studies have reported that *in vivo* genetic manipulation of PRMTs affected multiple aspects of skeletal muscle biology (see ^12^ for a detailed review). Notably: i) PRMT1 knockout (KO) resulted in a reduction in muscle mass and contractile force ^13^, ii) Co-activator associated arginine methyltransferase 1 (CARM1)/PRMT4 KO resulted in a reduction in muscle mass but also mitigated the progression of denervation-induced atrophy ^14^, iii) PRMT5 KO resulted in impaired myogenic differentiation and muscle regeneration ^15^ and, iv) PRMT7 KO resulted in a reduction in oxidative metabolism and exercise endurance ^16^. Furthermore, other studies have also implicated arginine methylation in metabolic diseases, including observations that PRMT1 KO in cultured myotubes reduced insulin-stimulated glucose uptake ^17^ and that PRMT7 content is reduced in mice with diabetes ^16^. Taken together, these emerging studies suggest that arginine methylation can act as an important mediator of skeletal muscle health and disease and that targeting methylarginines could be a viable therapeutic option to treat skeletal muscle pathologies. However, a comprehensive assessment of the regulation of the three distinct forms of methylarginine under physiological conditions of skeletal muscle health involving exercise and hypertrophy, and diseases involving metabolic disorders or atrophy has not been systematically performed.

Advances in mass spectrometry-based characterization of the arginine methylome have allowed for a greater understanding of the modified proteins and signaling pathways *in vivo*. In particular, work by the Nielsen group led to the identification of over 8,000 MMA sites in human HEK293 cells, which underpinned arginine methylation as one of the most common PTMs on par with protein phosphorylation and ubiquitination ^18^. Furthermore, the Ljubicic group revealed the remodeling of the mouse skeletal muscle arginine methylome in response to CARM1 deletion, including dynamic regulation of both monomethylation and dimethylation sites, which was accompanied by alterations in proteins involved in myofiber contraction, sarcomere organization and myogenesis ^19^. As such, research efforts utilizing quantitative mass spectrometry-based proteomics are vital in understanding the complex signaling cascades in health and disease. Nonetheless, identifying these regulated, discrete arginine methylation sites under (patho)physiological conditions in either human or mouse skeletal muscle has not been performed.

Thus, the first aim of this study was to investigate the physiological landscapes that regulate the three distinct variants of arginine methylation under settings of healthy skeletal muscle adaptations involving exercise and hypertrophic growth, and in diseases involving metabolism and atrophy. Our next aim was to identify skeletal muscle arginine methylation sites and quantify their changes under physiological settings. Our data revealed that all investigated atrophy models, including human ALS, resulted in changes in aDMA. Hence, we focused on the site-specific quantification of aDMA in skeletal muscle biopsies from individuals with ALS. In parallel, we also quantified the skeletal muscle proteome of human ALS, which, to our knowledge, represents the largest compendium of protein changes observed in a clinical cohort. These studies led us to investigate the effects of PRMT1 overexpression on aDMA, which increased fatigue resistance and functional recovery in adult mouse skeletal muscles.

## MATERIALS AND METHODS

### Mouse housing

All mouse experiments were approved by The University of Melbourne Animal Ethics Committee (#26527) or by the Victoria University Animal Ethics Committee (VUAEC #15/005), and fulfill the guidelines set out by the National Health and Medical Research Council of Australia for the care and use of animals for scientific purposes. Male C57BL/6J mice, *db/db*, *db/+*, FVB/N, SOD1^G37R^ and TDP-43^Q331K^ mice were housed at 22°C (±1°C) in groups of five/cage and maintained on a Standard Chow diet (Specialty Feeds, Glen Forest, WA, Australia, #SF15-049) with a 12-hr light/dark cycle and *ad libitum* access to food and water (see below for detailed experimental procedure). For the diet-induced insulin resistance model, male C57BL/6J mice were fed the standard chow diet or a high-fat diet (Specialty Feeds, #SF04-001) for 20 weeks.

### Ex vivo insulin-stimulated 2-deoxyglucose uptake

Mice were anesthetized with pentobarbitone sodium (270 mg/kg body weight) by intraperitoneal injection and transferred to a dissecting stage. Depth of anesthesia was assessed via the lack of leg and optical reflexes for at least 1 minute. After confirming anesthesia, skin from the hind legs was removed, the tendomuscular junction. Muscles were excised and incubated in Modified Krebs Buffer (MKB) (116 mM NaCl, 4.6 mM KCl, 1.16 mM KH2PO4, 25.3 mM NaHCO3, 2.5 mM CaCl2, 1.16 mM MgSO4) in a myograph (DMT, Hinnerup, Denmark, #820MS) at 30°C with constant gentle bubbling of 5% medical carbon dioxide in oxygen.

For 2-deoxyglucose uptake experiments, left and right soleus muscles were adjusted to resting tension (∼5 mN) and allowed to equilibrate for 10 min before being stimulated (left leg no insulin, right leg 100 nM insulin) for 30 min. During the last 10 min of stimulation, the buffer was replaced with MKB containing 0.375 μCi/mL 2-deoxy-d-[1,2-3H] glucose and 0.05 μCi/mL d-[14C] mannitol after which muscles were immediately immersed in ice-cold PBS to stop further glucose uptake. Each muscle was then excised of its sutures/tendons and frozen in liquid nitrogen for future lysis in 400 μL of 2% SDS via tip-probe sonication (QSonica, New Town, CT, USA). Measurement of radiolabeled 2-deoxyglucose was carried out in a Tri-Carb 4910 TR liquid scintillation analyzer (Perkin Elmer, Waltham, MA, USA) by adding 200 μL of muscle lysate to 4 mL scintillation fluid (Perkin Elmer, Ultima Gold). Glucose uptake rates were calculated as described previously ^20^.

### Diabetic mice model and experimental procedures

Male heterozygous *db/+* and homozygous *db/db* mice (Jackson Laboratory strain: BKS.Cg-*Dock7*^m^+/+ Lepr^db^/J) were maintained on a standard chow diet. At 12 weeks of age, mice were fasted for 4 hours, fasting blood glucose levels monitored (Roche Diagnostics, Castle Hill, NSW, Australia, Accu-Chek II glucometer), and mice gavaged orally with glucose (1 g/kg body weight), followed by assessment of blood glucose over 90 minutes. Quadriceps muscle was collected from animals fasted for four hours.

### Denervation-induced atrophy mice model and experimental procedures

For surgical denervation of the peroneal nerve, 8-week-old C57BL/6 male mice were placed under anaesthesia (2% isofluorane in O2) and then given a subcutaneous injection of carprofen for post-operative analgesia. A small incision was made in the skin distal to the knee to expose the peroneal nerve. Unilateral denervation was performed where the left peroneal nerve was exposed but not cut (Sham), while the right peroneal nerve was exposed, and a small piece (∼1l]mm) was excised (denervated). Following closure of the overlying incision and recovery of consciousness, mice were housed for 14 days post-denervation, as indicated.

### Synergistic ablation mice model and experimental procedures

Male FVB/N mice, aged 8–11 weeks, were purchased from the Animal Resources Center (ARC, Murdoch, WA) and housed at the Western Center for Health, Research and Education (Sunshine Hospital, St. Albans, Victoria, Australia). All surgeries were performed under isoflurane anesthesia, and following tissue extraction, mice were killed by cervical dislocation while still under anesthesia. To induce mechanical overload-induced muscle hypertrophy of the plantaris muscle, mice were subjected to synergist ablation (SA) surgery which involved the bilateral removal of the soleus (SOL) and distal half of the gastrocnemius muscle. Mice in control groups received sham surgery, for which an incision was made on the lower leg and then closed. After the surgical procedures, the incision was closed with Vetbond surgical glue (Henry Schein, Melville, NY, USA). Mice were allowed to recover for 3 or 7 d, after which mice were reanesthetized with isoflurane and the plantaris muscles were collected, weighed and snap frozen in liquid N_2_, and homogenized for western blot analysis as described previously ^21^.

### ALS mouse models and experimental procedures

The transgenic SOD1^G37R^ and TDP-43^Q331K^ mouse models of ALS used in this study were as previously described ^22,23^. In brief, SOD1^G37R^ mice develop rapidly progressive paralysis and die prematurely at around 28 weeks old, whereas TDP-43^Q331K^ mice develop a more slowly progressing, non-fatal ALS-like phenotype. SOD1^G37R^ mice and TDP-43^Q331K^ mice were sacrificed at 25 or 46 weeks for tissue collection, respectively. Non-transgenic littermates were used as controls. The procedures for killing, perfusion and tissue collection were all as previously described ^24^. Gastrocnemius muscle samples were snap-frozen on dry ice then stored at −80°C until analyzed.

### Human ALS participants and experimental procedures

Nine participants with ALS who met the revised El-Escorial criteria for ALS ^25^ were enrolled from the Royal Brisbane and Women’s Hospital (RBWH) motor neuron disease clinic for the collection of skeletal muscle biopsies. Nine healthy control participants were also enrolled. Control individuals were the spouses, friends or family members of ALS participants. For all participants, exclusion criteria were history of a metabolic condition (e.g. Hashimoto’s disease) and diabetes mellitus. Participant details are shown in **Table 1**. For ALS participants, the ALS Functional Rating Scale-Revised (ALSFRS-R) score and deltaFRS were obtained from clinical records. All participants provided written informed consent; participant consent was obtained according to the Declaration of Helsinki. Work performed in this study was approved by the RBWH and University of Queensland human research ethics committees.

**Table 1:** Summary of participant characteristics.

|  | ALS (n = 9) | Control (n = 9) | p-value |
| --- | --- | --- | --- |
| Age (years) | 56.9 ± 6.49 | 61.5 ± 10.1 | 0.34 |
| Sex (Female) | 3 (33.3%) | 2 (22.2%) | 0.60 |
| BMI (kg/m<sup>2</sup>) | 25.0 ± 4.43 | 24.8 ± 2.17 | 0.55 |
| % Fat | 31.3 ± 12.6 | 27.8 ± 7.81 | 0.60 |
| Fat-free mass (kg) | 51.0 ± 10.3 | 54.9 ± 9.79 | 0.26 |
| FVC (% predicted) | 84.4 ± 7.7 |  |  |
| SNIP (% predicted) | 52.5 ± 7.0 |  |  |
| ALSFRS-R | 35.8 ± 2.7 |  |  |
| ΔFRS | -0.5 ± 0.12 |  |  |

Measured age (years), sex (Female), body mass index (BMI; kg/m<sup>2</sup>), % fat mass, fat mass (kg), lean mass (kg), forced vital capacity (FVC; % predicted) and sniff nasal inspiratory pressure (SNIP; % predicted) in ALS (n = 9) and control participants (n = 9). The ALS functional rating scale (ALSFRS-R) and change in ALSFRS-R since time of symptom onset (ΔFRS, points/month) were assessed at time of sample collection. FVC, SNIP, ALSFRS-R and ΔFRS were not tested in control participants. Data depicted as mean +/- SD; α = 0.05; Data analyzed using the Mann-Whitney U-test, except for categorical values, where data analyzed using the Chi-square test.

#### Assessment of body composition and energy expenditure

Body composition (fat mass and fat-free mass) was determined by whole-body air displacement plethysmography using the BodPod system (Cosmed, Albano Laziale, Rome, Italy) ^26^. Values of fat mass and fat-free mass were used to predict resting energy expenditure) ^27^. Energy expenditure in fasting and alert ALS and control participants at rest (i.e. resting energy expenditure) was measured by indirect calorimetry using a Quark RM respirometer (Cosmed) as we have done previously ^27^.

#### Muscle biopsy

Muscle biopsies were collected from the *vastus lateralis* of one leg. Lignocaine (1%; 5ml) was injected to anaesthetize the skin, underlying fat and muscle tissue. A 10mm incision was made and advanced through the fascia of the muscle. A ∼200mg sample of muscle was collected using a sterile 6mm hollow Bergstrom biopsy needle (Pelomi, Albertslund, Denmark) modified for suction ^28^and placed in holding media for transport to the laboratory (Thermo Fisher Scientific, Waltham, MA, USA, DMEM/F12 with 0.5% gentamicin). Samples were washed in PBS, frozen on dry ice, and stored in liquid nitrogen.

### Human exercise participants and experimental procedures

The human exercise interventions, biopsy collection and lysates for western blotting were generated as described previously ^29^.

### Lysate preparation

Tissue was lysed by tip-probe sonication in 6M guanidine HCl (Sigma-Aldrich, St. Louis, MO, USA, #G4505), 100mM Tris pH 8.5 containing 10mM Tris(2-carboxyethyl)phosphine (Sigma-Aldrich, #75259) and 40mM 2-Chloroacetamide (Sigma-Aldrich, #22790). The lysate was then heated at 95°C for 5 minutes and centrifuged at 18,000 g for 20 minutes at 4°C. The supernatant was diluted 1:1 with LC-MS/MS water and precipitated overnight in a final concentration of 80% acetone at −30°C. The lysate was centrifuged at 4,000 g for 5 minutes at 4°C and the supernatant was disregarded. The protein pellet was washed with 80% ice-cold acetone and centrifuged at 4,000 g for 5 minutes at 4°C, then resuspended in digestion buffer (10% trifluoroethanol (Thermo Fisher Scientific, #96924) in 100 mM HEPES pH 7.5). Protein was quantified using a BCA protein assay (Thermo Fisher Scientific, #23325) and normalized to 2µg/µl containing Laemelli buffer (BioRad, Hercules, CA, USA, #1610747) for western blotting. For proteomics, 1mg of protein was normalized to 200μl of digestion buffer, then digested in sequencing grade trypsin (Sigma-Aldrich, #T6567) and LysC (Wako, Chuo-Ku, Osaka, Japan, #129-02541) (1:100 enzyme:substrate ratio) overnight at 37°C with shaking at 2000 rpm.

### Peptide purification

Peptides were acidified by Trifluoroacetic acid (TFA) (Thermo Fisher Scientific, #FSBA116) to a final concentration of 1% TFA and centrifuged at 16,000 g for 10 minutes at room temperature. Peptides were desalted using reversed-phase chromatography with HLB-SPE (30mg) columns (Waters, Milford, MA, USA, WAT094225) and washed with 0.1% TFA, then eluted with 80% acetonitrile, followed by 0.1% TFA. Peptides were vacuum-dried at 45°C for at least 45 minutes.

### Arginine methylpeptide enrichment

Arginine dimethylated peptides were enriched by immunoprecipitation (IP) using Asymmetric Di-Methyl Arginine (adme-R) antibodies coupled to magnetic immunoaffinity beads (Cell Signalling Technology, Danveres, MA, USA, #18303). Peptides were resuspended in immunoprecipitation buffer (50mM MOPS, 10mM NaH_2_PO_4_, 50mM NaCl, pH 7.5), and adjusted to a pH of 7.5 with 5M NaOH, then centrifuged at 16,000 g for 5 minutes at 4°C. The beads were washed twice with IP buffer then transferred to the peptide solution and incubated overnight at 4°C on a rotator. Dimethylpeptides were washed three times with IP buffer, followed by LC-MS/MS water, then dried. Dimethylpeptides were then eluted with 0.2% TFA for 5 minutes at 10°C with shaking at 1200 rpm, and beads filtered using in-house packed C8 tips (3M, Saint Paul, MN, USA, #11913614). The purified peptides were resuspended in 200mM HEPES, pH 7.4, for 5 minutes with shaking at 2000 RPM, then labelled with TMTpro-16plex (Thermo Fisher Scientific, #A44520), followed by 1 hour incubation at room temperature. The reaction was deacylated to a final concentration of 0.3% hydroxylamine then quenched with 1% TFA. All 16 samples were pooled together and purified through in-house packed SDB-RPS (Sigma-Aldrich, #66886-U) tip, and washed with 99% isopropanol, 1% TFA followed by 5% acetonitrile, 0.2% TFA, then eluted with 80% acetonitrile, 5% NH_4_OH. Sample was resuspended overnight in 2% acetonitrile, 0.1% TFA and stored at −80°C, then fractionated using a HpH C18 column into 12 concatenated fractions as previously described ^30^.

### LC-MS/MS data acquisition

Peptides for total proteomic analysis were analyzed on a Dionex 3500 nanoHPLC coupled to an Orbitrap Eclipse mass spectrometer (Thermo Fisher Scientific) via electrospray ionisation in positive mode with 1.9 kV at 275 °C and RF set to 30%. Separation was achieved on a 50 cm × 75 µm column packed with C18AQ (Dr Maisch, Ammerbuch, Germany, 1.9 µm) (PepSep, Marslev, Denmark) over 60 minutes at a flow rate of 300 nL/min. The peptides were eluted over a linear gradient of 3–25% Buffer B (Buffer A: 0.1% formic acid [FA]; Buffer B: 90% v/v acetonitrile, 0.1% v/v FA) and the column was maintained at 50 °C. The instrument was operated in data-independent acquisition (DIA) mode with an MS1 spectrum acquired over the mass range 350–1,400 *m/z* (120,000 resolution, 1 x 10^6^ automatic gain control (AGC) and 50 ms maximum injection time) followed by MS/MS analysis of 50 x 13.7 *m/z* isolation windows over the mass range of 360.5 – 1,033.5 *m/z* via HCD fragmentation mode and detection in the orbitrap (30,000 resolution, 1 × 10^6^ AGC, 55 ms maximum injection time; 30% normalized collision energy). Fractionated TMT labelled peptides following dimethyl arginine enrichment were analyzed on the identical chromatography conditions as described above except separation was achieved over 45 minutes. The instrument was operated in data-dependent acquisition (DDA) mode with an MS1 spectrum acquired over the mass range 350–1,550 *m/z* (120,000 resolution, 1 x 10^6^ automatic gain control (AGC) and 50 ms maximum injection time) followed by MS/MS analysis via HCD fragmentation mode and detection in the orbitrap (30,000 resolution, 1 × 10^5^ AGC, 59 ms maximum injection time; 35% normalized collision energy, Enhanced Resolution Mode for TMTpro reagents, 3 second cycle time, 1.3 *m/z* isolation, dynamic exclusion 30 seconds).

### LC-MS/MS data processing

Data for total proteomic acquisition (LFQ-DIA) were searched against the UniProt human database (March 2023) with Spectronaut (v17.6.230428.55965) using default parameters with peptide spectral matches (PSM), peptide and protein false discovery rate (FDR) set to 1%. All data were searched with oxidation of methionine and N-terminal protein acetylation set as the variable modification and carbamidomethylation of cysteine set as the fixed modification. Peptide quantification was carried out at MS2 level using 3-6 fragment ions, with automatic interference fragment ion removal as previously described ^31^. The MS1 mass tolerance was set to 20 ppm, while the mass tolerance for MS/MS fragments was set to 0.02 Da. Dynamic mass MS1 and MS2 mass tolerance was enabled, and retention time calibration was accomplished using local (non-linear) regression. A dynamic extracted ion chromatogram window size was performed. Data for dimethyl proteomic acquisition (TMT-DDA) were searched against the UniProt human database (March 2023) with SequestHT and Percolator ^32^ in Proteome Discoverer (v2.5.0.4) and filtered to 1% FDR at the PSM, peptide and protein level. All data were searched with oxidation of methionine, N-terminal protein acetylation, and methylation and dimethylation of arginine set as the variable modification. Carbamidomethylation of cysteine, and TMTpro of lysine and peptide N-terminus were set as fixed modification. The MS1 mass tolerance was set to 10 ppm, while the mass tolerance for MS/MS fragments was set to 0.02 Da. Quantification was performed with the reporter ion quantification node for TMT quantification in Proteome Discoverer. TMT precision was set to 20 ppm and corrected for isotopic impurities. Only spectra with < 50% co-isolation interference were used for quantification with an average signal-to-noise filter of > 10. Localisation of methylation sites was performed with PhosphoRS ^33^. Data were processed with Perseus ^34^ with log2-transformation and normalization by subtracting the median of each sample. Statistical analysis was performed using student’s t-tests and correcting for multiple hypothesis testing using Benjamini Hochberg with significance set to 5% FDR. Pathway enrichment analysis was performed with Metascape ^35^. GSEA against ChEA3 ^36^ was performed in TeaProt ^37^. Protein:protein interaction analysis was analysis was performed on STRING (v12.0). MCODE analysis was performed on Cytoscape (v3.10.0).

### Western blotting

Proteins were separated on NuPAGE 4-12% Bis-Tris protein gels (Thermoscientific) in MOPS SDS running buffer at 145V for 60 minutes in room temperature. Proteins were then transferred to PVDF membranes (Millipore, Burlington, MA, USA, #IPFL00010) in NuPAGE transfer buffer at 20V for 60 minutes at room temperature. Membranes were blocked with 5% BSA in Tris-buffered saline containing 0.1% Tween-20 (TBST) for at least 60 minutes at room temperature with gentle shaking. Membranes were then incubated overnight at 4°C with gentle shaking in primary antibody containing 5% BSA, 0.02% NaN3 in TBST. The antibodies used were Mono-methyl Arginine MultiMab (1:500) (Cell Signalling Technology, #8015), Asymmetric Di-methyl Arginine MultiMab (1:500) (Cell Signalling Technology, #13522), Symmetric Di-methyl Arginine MultiMab (Cell Signalling Technology, #13222) (1:500), PRMT1 (1:1000) (Cell Signalling Technology, #2449), Phospho-S6 Ribosomal Protein (Ser235/236) (1:1000) (Cell Signalling Technology, #4856), Phospho-Akt (Ser473) (1:1000) (Cell Signalling Technology, #4060) and Akt (1:1000) (Cell Signalling Technology, #9272). Membranes were incubated with HRP-secondary antibody in 5% BSA in TBST for at least 60 minutes at room temperature with gentle shaking. Protein was visualized with Immobilon Western Chemiluminescent HRP Substrate (Millipore, #WBKLS0500) and imaged on a ChemiDoc (BioRad). Densitometry was performed in Image J ^38^.

### PRMT1 myoAAV production

MyoAAV ^39^ vectors were produced by the Vector and Genome Engineering Facility (VGEF) at Children’s Medical Research Institute (CMRI, Westmead, NSW, Australia). Vectors were produced by standard transient transfection of 5 x 15cm plates of HEK293 cells using PEI (polyethylenimine) (PolyPlus, Illkirch-Graffenstaden, France, #115-100) with a 1:1:2 molar ratio of pTransgene:pRep2CapXHelper:pAd5Helper. Vectors were purified using iodixanol gradient ultracentrifugation as previously described ^40^. Amicon Ultra-4 Centrifuge Filter Units (Ultracel-100 kDa membrane) (Millipore, #UFC810024) were used to perform buffer exchange (Phosphate-buffered saline [PBS] [Gibco, #14190], 50 mM NaCl [Sigma-Aldrich, #S5150-1L], 0.001 %, Pluronic F68 [v/v] [Gibco, #24040]) and the final concentration step. Iodixanol-purified AAVs were quantified using droplet digital PCR (ddPCR) (BioRad, Berkeley) using QX200 ddPCR EvaGreen Supermix (Bio-Rad, 1864034) with eGFP primers (5’ TCAAGATCCGCCACAACATC and 5’ TTCTCGTTGGGGTCTTTGCT). All cell stocks were regularly checked for absence of mycoplasma with the Mycoplasma Detection Kit (Jena Bioscience, Jena, Thüringen, Germany, #PP-401).

### PRMT1 myoAAV injection

7 weeks old C57BL/6J male mice (WEHI, Parkville, VIC, Australia) (n = 10) were anaesthetised with 4% isoflurane in oxygen at 1 L/min, then transferred to a heated dissecting microscope stage with an isoflurane inhalation nose piece (2% in oxygen at 1 L/min). Depth of anaesthesia was assessed by the lack of hindlimb reflexes for at least 1 minute and the position of the head was checked to ensure normal breathing. The surface of the hindlimb was sterilised with 80% ethanol. TA muscle was injected with 1−10^9^ vector genome/30 µl of PRMT1 myoAAV or eGFP myoAAV using a 32G needle. Mice were returned to their cages, and their body weights were monitored daily for the first 3 days and then weekly.

### Muscle dissection

Mice were anesthetized as outlined above. After confirming anesthesia, skin from the hindlimbs was removed, and the EDL and TA muscles were excised. EDL muscles were weighted then subjected to ex-vivo muscle function testing. TA muscles were weighted then immediately snaped frozen in liquid nitrogen, and later used for western blotting and proteomics.

### Ex-vivo muscle function testing

EDL muscles were sutured at the proximal and distal ends of the tendomuscular junction using 4-0 suture, then incubated in modified Krebs buffer (116 mM NaCl, 4.6 mM KCl, 1.16 mM KH2PO4, 25.3 mM NaHCO3, 2.5 mM CaCl2, 1.16 mM MgSO4, 0.1% BSA, 2mM pyruvate, 11mM glucose) in a myograph (DMT, #820MS), with gentle bubbling of 5% carbogen at 30°C. A constant voltage stimulator (DMT, CS4) was used to deliver 0.2ms supramaximal pulses (26V) via stimulation electrodes (DMT, 300415) placed over the mid-belly of the muscle. Optimal muscle length was determined by delivering successive twitch stimulation, with at least 30 seconds rest by incrementally stretching the muscle until maximum twitch force was obtained. Force-frequency relationship was determined by stimulating muscles with incrementally increasing frequencies (10 to 200Hz, 350ms duration), with at least 2 minutes rest between stimulation. Muscle fatigue was assessed by maximally stimulating the muscle once every 4 seconds for 4 minutes. Recovery was assessed by peak tetanic force production by maximally stimulating the muscle 5, 10 and 15 minutes after fatigue. Force values were normalized to muscle cross-sectional area by diving muscle mass by the product of muscle length and muscle density (1.06 mg/mm^3^). Force recordings were digitized using a PowerLab 8/35 unit (ADInstruments, Dunedin, NZ) and analyzed using the Peak Parameters module in LabChart Pro (v8.1.16, ADInstruments).

### Statistical analysis

Statistics for SA, exercise, HFD, *db/db*, denervation and ALS results were performed in Graph-Pad Prism (Version 9.5.1). Paired t-test was used to analyze data related to SA experiments, including muscle mass and phospho/methyl blots. Two-way ANOVA was used to analyze data related to exercise experiments including western blots. Two-way ANOVA was used to analyze data related to HFD experiments, including 2DG uptake and phospho/methyl blots (unpaired t-test was used for 2DG and pAKT fold change). Unpaired t-test was used to analyze data related to *db/db* experiments, including fasting blood glucose, GTT and western blots. Paired t-test was used to analyze results relating to denervation-induced atrophy, including western blots. Unpaired t-test were used to analyze results relating to ALS (SOD1^G37R^, TDP-43^Q331K^, and human). Mann-Whitney U-test were used to analyze ALS versus control participant characteristics except for categorical values, where the Chi-square test was used. All mentions of sample size (n) refer to biological replicates.

## List of antibodies

MMA: RRID:AB_2799401

aDMA: RRID:AB_2665370

sDMA: RRID:AB_2714013

PRMT1: RRID:AB_2237696

p-RPS6(S235/6): RRID:AB_2181037

p-AKT(S473) RRID:AB_2315049

AKT: RRID:AB_329827

## RESULTS

### Global quantification of arginine methylation in models of skeletal muscle health

We first quantified the global regulation of arginine methylation in two models representative of physiological adaptations in healthy skeletal muscle: 1) hypertrophic growth induced by synergistic ablation (SA) in mice and 2) various acute exercise modalities in humans.

SA was induced by the bilateral removal of the soleus muscle and distal half of the gastrocnemius muscle, leading to functional overload and rapid hypertrophy of the plantaris muscles ^41^. The mice recovered for 3 or 7 days (n = 3; each), at which point their plantaris muscles were harvested. SA led to a significant increase in plantaris muscle mass at day 7 (**Figure 1A**), and western blot confirmed activation of hypertrophic signaling with an increase in pS235/6 on RPS6, a downstream substrate of mTORC1/P70S6K (**Figure 1B**). MMA and aDMA levels did not differ between groups after 3 days but were increased following 7 days of SA compared to the sham-treated muscle (**Figure 1C**).

**Figure 1.**
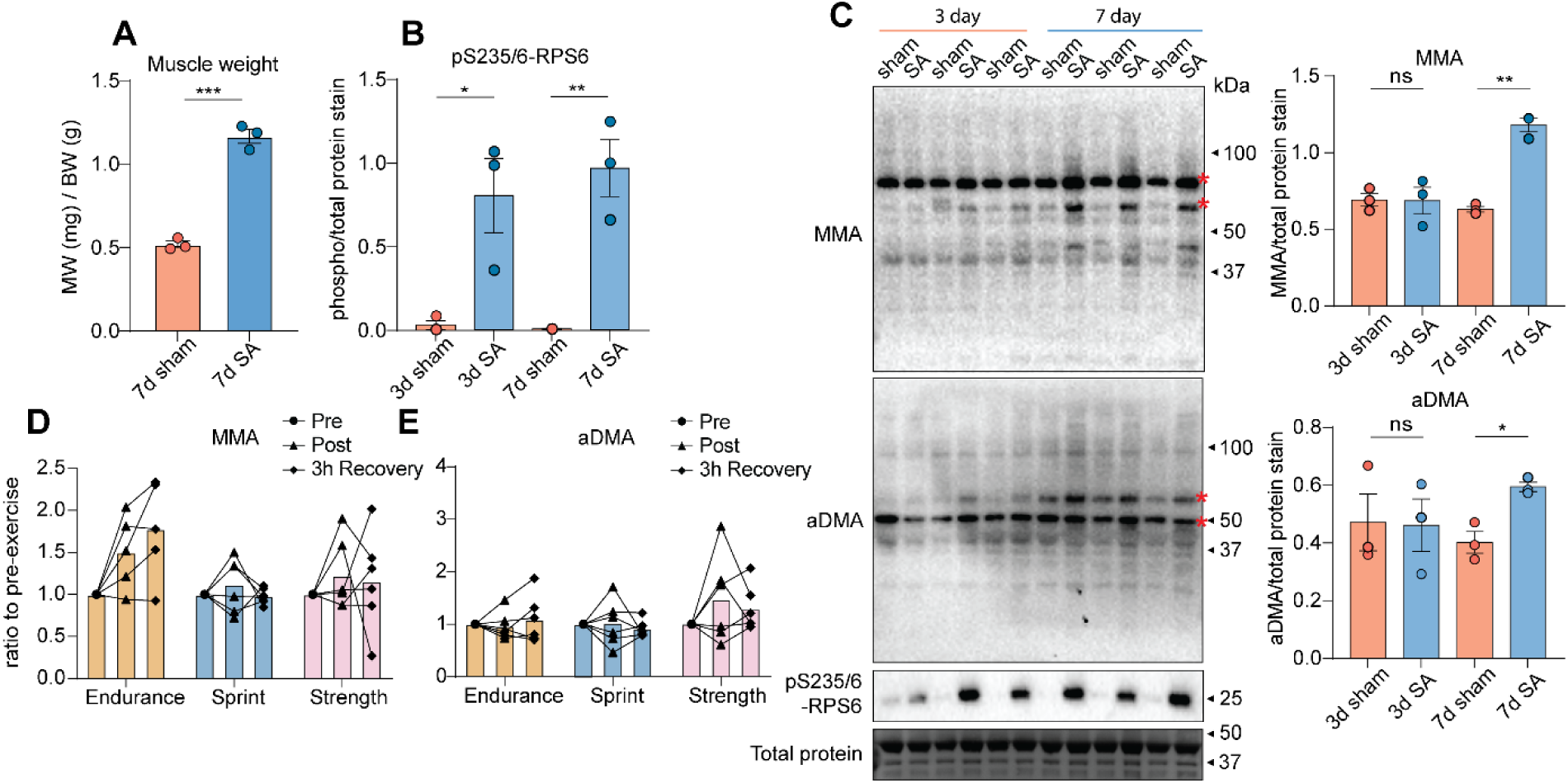
Global quantification of arginine methylation in models of skeletal muscle health. **A**). Plantaris muscle weight (MW)/body weight (BW) ratio (n = 5) of mice after 7-day sham or 7-day synergistic ablation (SA). **B**). Quantification of phosphorylated ribosomal protein S6 (p-RPS6) in plantaris muscles following 3-day and 7-day sham or SA in mice (n = 3; each). **C**). Western blot and quantification of monomethyl arginine (MMA) and asymmetric dimethyl arginine (aDMA) in plantaris muscles following 3-day and 7-day sham or SA in mice (n = 3; each). Quantification of MMA (**D**) and aDMA (**E**) levels in human exercise muscle biopsies (n = 6) taken pre-exercise, immediately post-exercise, and 3 hours into recovery across three different exercise modalities (endurance, sprint and strength). Values normalized to pre-exercise levels. Data are represented as mean ± SEM; *p < 0.05, **p < 0.01, ***p < 0.001; α = 0.05; Red asterisk denotes the band(s) in which quantification was performed (summed densitometry); (A-C) Unpaired t-test; (D-E) One-way ANOVA with Tukey post hoc test.

Next, we sought to understand the regulation of arginine methylation during acute exercise in humans. Six healthy untrained men performed acute bouts of endurance (90 minutes, 60% of VO_2_ max), sprint (3 x 30 seconds all-out cycling) and resistance (6 sets of 10 repetitions maximum knee extension) exercise on separate days as previously described ^29^. Muscle biopsies from *m. vastus lateralis* were obtained before, immediately after exercise and 3 hours post-recovery. Western-blotting/phosphoproteomics was previously performed to confirm robust and acute activation of various phosphorylation-based exercise signaling pathways in the different modalities in all participants ^29^. Western blot analysis showed variable levels of global MMA and aDMA between the participants and revealed no significant differences across the three exercise modalities (**Figure 1D-E**). Although some participants displayed subtle changes in arginine methylation, protein phosphorylation analysis of muscle biopsies from the same 6 participants revealed robust changes, such as a >10-fold increase in the phosphorylation of S80 on ACC ^29^. The stark contrast in the degree of methylation versus phosphorylation suggests that arginine methylation does not play a major role in acute exercise response. Taken together, although we observed a minor increase in arginine methylation following SA-induced hypertrophy, our data suggest that under physiological settings that promote muscle health, such as growth and acute exercise, this PTM does not appear robustly regulated.

### Skeletal muscle arginine methylation is not regulated under settings of insulin resistance or T2D

We next investigated the regulation of arginine methylation in two mouse models of metabolic disease: 1) insulin resistance induced by 20 weeks of high-fat feeding (HFD), and 2) T2D induced by mutation of the leptin receptor (Lepr^db/db^). HFD feeding significantly blunted insulin-stimulated glucose uptake (**Figure 2A-B**) and attenuated insulin-stimulated phosphorylation of S473 on Akt **(Figure 2C-D**), confirming skeletal muscle insulin resistance. Western blot analysis revealed no significant changes in the levels of MMA, aDMA and sDMA between chow and HFD mice under both basal conditions and during insulin stimulation (**Figure 2E-H**).

**Figure 2.**
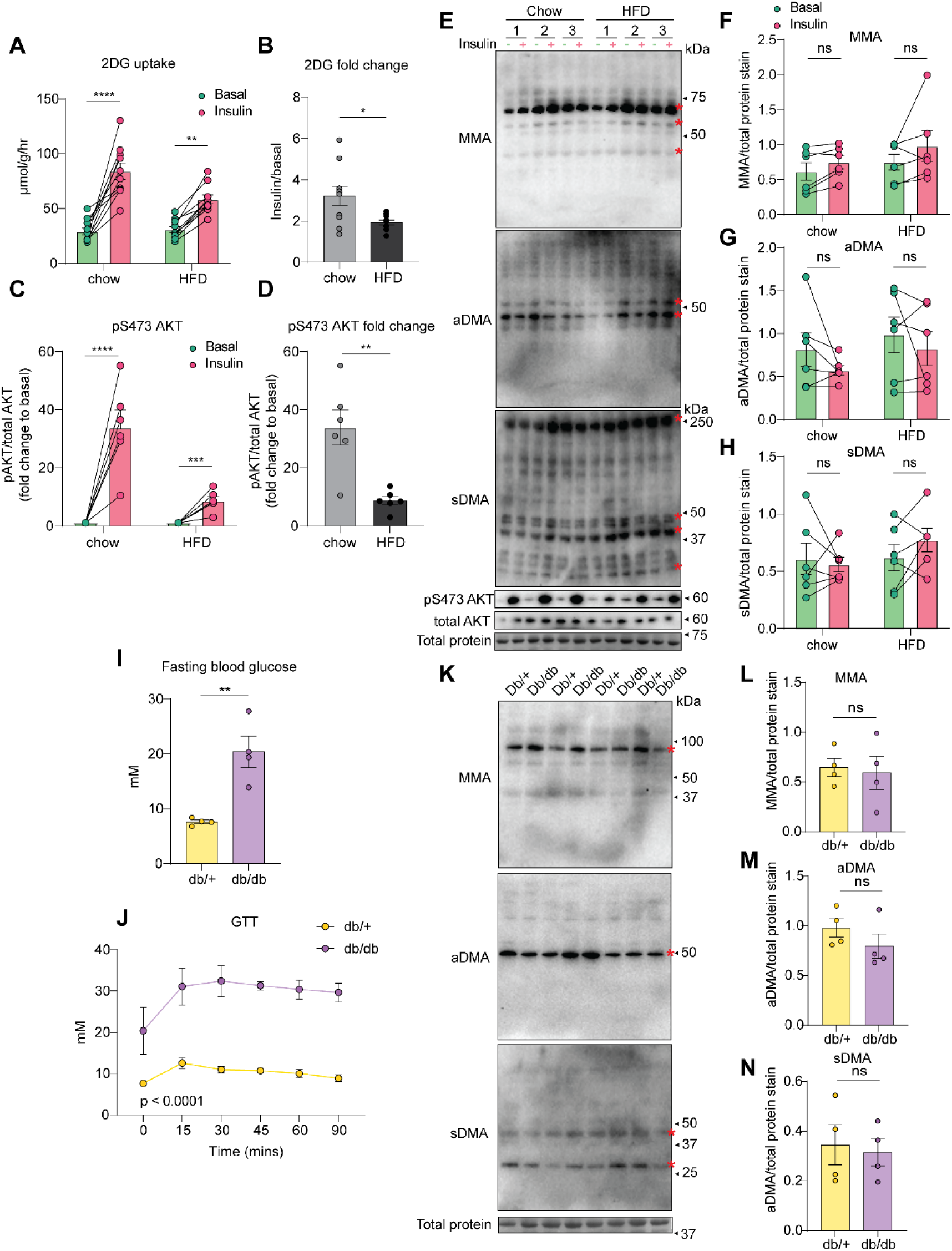
Global quantification of arginine methylation in skeletal muscle of insulin resistance and T2D mice. **A**). *Ex vivo* 2-DG uptake (µmmol/g/hr) in soleus muscles of basal left leg and insulin-stimulated right leg in chow and high-fat diet mice (n = 10; insulin = 100 mM; 30 min). **B**). 2-DG fold change (insulin/basal) in chow and high-fat diet mice (n = 10). **C**). S473 phosphorylation on AKT in soleus muscles of basal left leg and insulin-stimulated right leg in chow and high fat diet mice (n = 3), normalized to basal level. **D**). pS473 phosphorylation on AKT fold change (insulin/basal) in chow and insulin resistance mice (n = 3)f. Western blot (**E**) and quantification of MMA (**F**), aDMA (**G**), and sDMA (**H**) in left basal and right insulin-stimulated soleus muscle of chow and insulin resistance mice (n = 6). **I**) Fasting blood glucose levels (mM) in diabetic *db/db* and control *db/+* mice (n = 4) **J**). Measurements of blood glucose levels (mM) at 0, 15, 30, 35, 60 and 90 minutes in diabetic *db/db* and control *db/+* mice subjected to a glucose tolerance test (n = 4). Western blot (**K**) and quantification of MMA (**L**), aDMA (**M**), and sDMA (**N**) in quadriceps muscles of diabetic *db/db* and control *db/+* mice (n = 4). Quantified MMA, aDMA and sDMA levels were normalized against total protein by densitometry. Data are represented as mean ± SEM; ns = non-significant; α = 0.05; Red asterisk denotes the band(s) in which quantification was performed (summed densitometry); (A, C, F-H) Two-way ANOVA; (B, D, I-J, L-N) Unpaired t-test.

We next investigated the regulation of arginine methylation in T2D using the *db/db* mouse model. *db/db* mice are homozygous for a mutation of the leptin receptor, a major sensor of energy homeostasis. As a result, these mice have defective leptin signaling and develop hyperphagia and morbid obesity ^42^. Mice were hyperglycemic (**Figure 2I**) and profoundly glucose intolerant (**Figure 2J**) as compared to heterozygous littermates (*db/+*). Western blot analysis revealed no significant differences in all three methylarginines: MMA, aDMA and sDMA, between *db/db* and *db/+* control mice (**Figure 2K-N**).

Taken together, these data indicate that arginine methylation in the skeletal muscles of mice is not altered in models of insulin resistance.

### Asymmetric dimethyl arginine (aDMA) is regulated in atrophy

We next investigated the regulation of skeletal muscle arginine methylation in settings of atrophy using a model of denervation-induced atrophy in mice, two transgenic mouse models (SOD1^G37R^ and TDP-43^Q331K^) of ALS, and in skeletal muscle biopsies from patients with ALS.

Denervation-induced atrophy was induced by unilateral peroneal denervation of the right hindlimb and sham surgery in the contralateral left hindlimb. After 14 days, tibialis Anterior (TA) muscles were harvested, and as expected, there was profound atrophy in the denervated limb as previously described ^43^. Here, we focused on western blot analysis of aDMA since pilot studies showed no differences in MMA and sDMA (data not shown). Western blot analysis revealed fascinating changes in methylation, with some bands not changing (e.g ∼45 kDa) and some undergoing significant increases (e.g ∼60 kDa) in aDMA levels in the denervated muscles compared to the sham control (**Figure 3A**), suggesting complex and site-specific regulation in methylation of discrete proteins.

**Figure 3.**
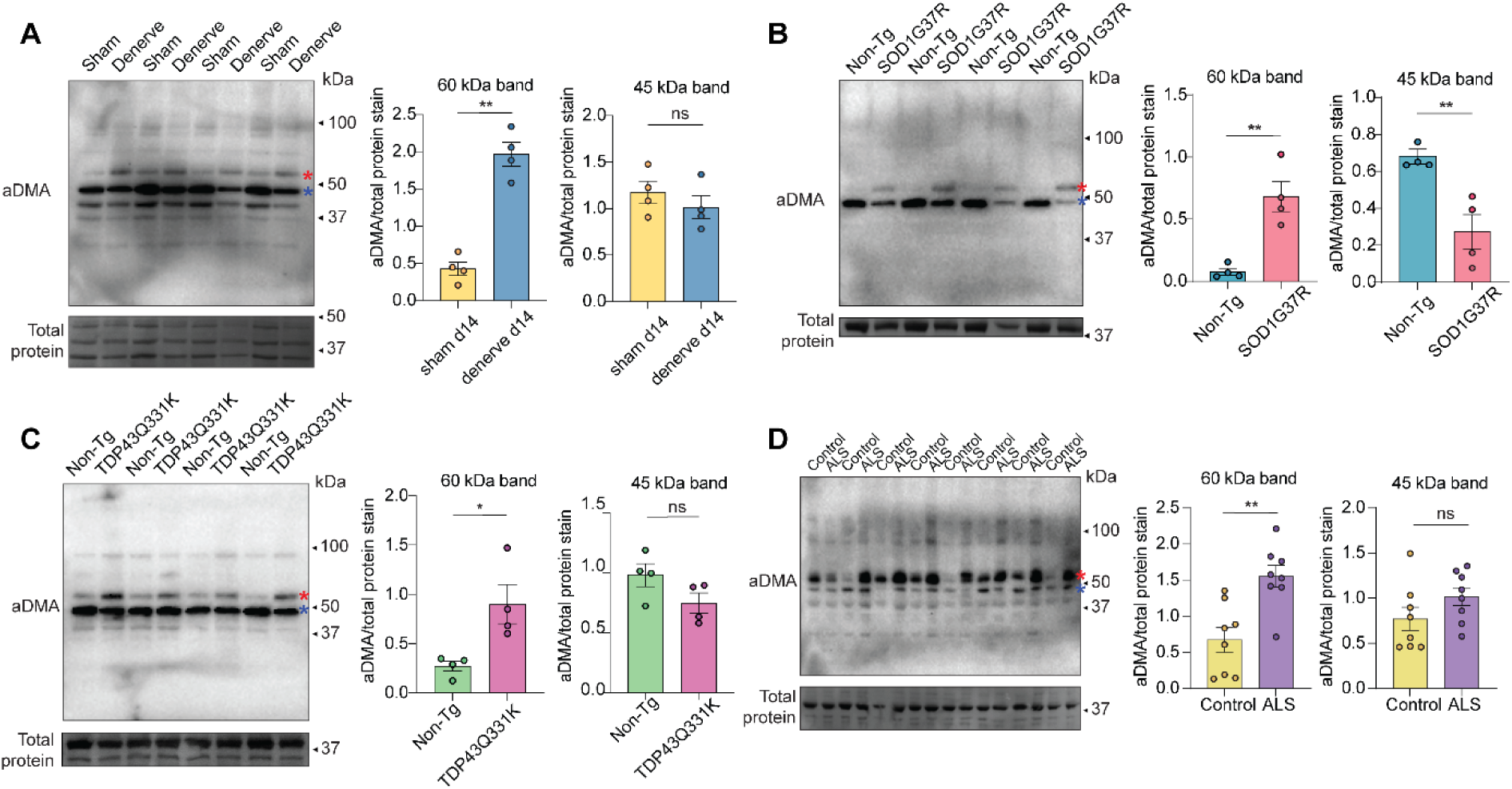
Global quantification of asymmetric dimethyl arginine (aDMA) in atrophic skeletal muscles. **A**) Western blot and quantification of asymmetrical dimethyl arginine (aDMA) in right TA muscle with denervation-induced atrophy and the left TA sham-treated controls in mice (n = 4). **B**) Western blot and quantification of aDMA in gastrocnemius muscles of 25 weeks old ALS mice induced by transgenic superoxide dismutase 1 (SOD1^G37R^) mutation and non-transgenic controls (n = 4). **C**) Western blot and quantification of aDMA in gastrocnemius muscles of 46 weeks old ALS mice induced by TAR DNA binding protein 43 (TDP-43^Q331K^) mutation and non-transgenic controls (n = 4). **D**) Western blot and quantification of aDMA in muscle biopsies from humans with ALS and aged-match controls (n = 8). All quantified aDMA levels were normalized to total protein. Data are represented as mean ± SEM; ns = non-significant; α = 0.05; (A-C) Red asterisk denotes the ∼60kDa band in which densitometry was performed; (A-C) Blue asterisk denotes the ∼45kDa band in which densitometry was performed. (D) Red asterisk denotes the ∼60kDa band which densitometry was performed. (A) Paired t-test; (B-D) Unpaired t-test.

Next, we investigated the regulation of arginine methylation in two transgenic mouse models of ALS - superoxide dismutase 1 (SOD1^G37R^) and TAR DNA binding protein 43 (TDP-43^Q331K^). These mouse models develop extensive neuromuscular atrophy and progressive muscle weakness ^22,23^, thus developing phenotypes reminiscent of ALS in humans, though the TDP-43^Q331K^ model is often considered less clinically severe. Western blot analysis revealed significant increases in protein aDMA levels in both SOD1 (**Figure 3B**) and TDP-43 mice (**Figure 3C**), at ∼60 kDa, compared to non-transgenic control mice. Interestingly, the protein band between ∼45 kDa had significant decreases in aDMA levels in the SOD1 mutant but not the TDP-43 mutant. Finally, we interrogated changes in arginine methylation in ALS *vastus lateralis* biopsies obtained from 6 males and 2 females with ALS, and 8 age/sex match individuals without ALS (see **Table 1** and **S1** for participant characteristics). Western blot analysis revealed significant increases in aDMA levels in ALS compared to control participants at ∼60 kDa, consistent with the results observed from the SOD1 and TDP-43 mice mutants (**Figure 3D**). There was considerable variation in immunoreactivity in the band at ∼45 kDa, along with a trend for increasing aDMA in several proteins at ∼100 kDa.

Taken together, our data reveal that aDMA is remodeled on specific proteins, likely in a site-specific manner, in mouse models of atrophy induced by the loss of innervation. These changes were also observed in skeletal muscle biopsies from ALS participants.

### Quantification of the skeletal muscle proteome and asymmetric dimethylome in human ALS

Our investigation into the global changes of arginine methylation in skeletal muscle health and disease has revealed the regulation of asymmetric dimethylation as a prominent feature of atrophy induced by loss of innervation. Thus, we next performed an integrated proteomic analysis to identify and quantify the exact proteins containing these regulated dimethylation sites. Here, we focused on characterizing the arginine dimethylome in skeletal muscle from ALS participants and age/sex match controls. **Table 1** presents the summary participant characteristics used in this study, revealing non-significant differences across all measurements between ALS and control participants.

Quantification of the skeletal muscle proteome was performed by analyzing Trypsin/LysC digested proteins from the *vastus lateralis* muscles of 9 patients with ALS and 9 age-match controls using 1D-nanoLC-MS/MS (**Figure 4A**). We identified 3,686 unique protein groups, with 3,140 proteins quantified in all 9 ALS samples and 2,946 proteins quantified in all 18 samples (**Figure 4B**) (**Table S1**). Principle component analysis and unsupervised hierarchical clustering revealed robust segregation of the proteome between ALS and control participants (**Figure 4C** and **D**). 572 proteins were significantly upregulated, and 221 proteins were significantly downregulated in ALS, making for a total of 793 differentially regulated proteins (**Figure 4E)**. The most significantly upregulated protein was actin-binding protein – MTSS I-BAR domain containing 1 (MTSS1), and the most significantly downregulated protein was cytokine receptor-like factor 3 (CRLF3), a negative regulator of cell cycle progression. We identified significant upregulation of FUS (fused in sarcoma) that is often mutated in cases of ALS ^44^. Surprisingly, we did not detect C9ORF72, which was recently identified as the most common genetic cause of ALS, accounting for more than 40% of familial and 8% of sporadic cases ^45^. To gain an overview of the regulated cellular pathways, enrichment analysis was performed against the Kyoto Encyclopedia of Genes and Genomes (KEGG) database (**Table S2**) ^46^. Significantly upregulated pathways in ALS were associated with pre-mRNA splicing, protein processing in the ER, phagosome and ribosomal proteins, while pathways involved in central carbon/amino acid/carbohydrate metabolism, motor proteins and calcium signalling were the most downregulated (**Figure 4F**) (**Table S2**). We next estimated the activity of transcription factors by performing a gene set enrichment analysis of target genes with the CHEA3 database ^36^. ALS was associated with elevated activity of TEAD4, TCF12, IKZF1 and GATA3 which, to our knowledge, have not previously been associated with ALS pathology (**Figure 4G**). Finally, we investigated the co-regulation of protein complexes retrieved from the STRINGdb ^47^ (**Figure 4H**). Consistent with pathway analysis, we observed an increase in the ribosome and spliceome complex. We also observed an increase in the Dynactin complex, the AP-2 complex and proteins forming the Tight Junction. ALS reduced the abundance of F-Actin capping enzymes, proteins forming the Z-Disk, the Myosin Complex, proteins contributing to the mitochondrial permeability transition pore (mPTP), enzymes regulating coenzyme Q biosynthesis, several major calcium transporters and proteins forming a SCF-like Elongin-Cullin-SOCS-box protein (ECS) E3 ubiquitin-protein ligase complex.

**Figure 4.**
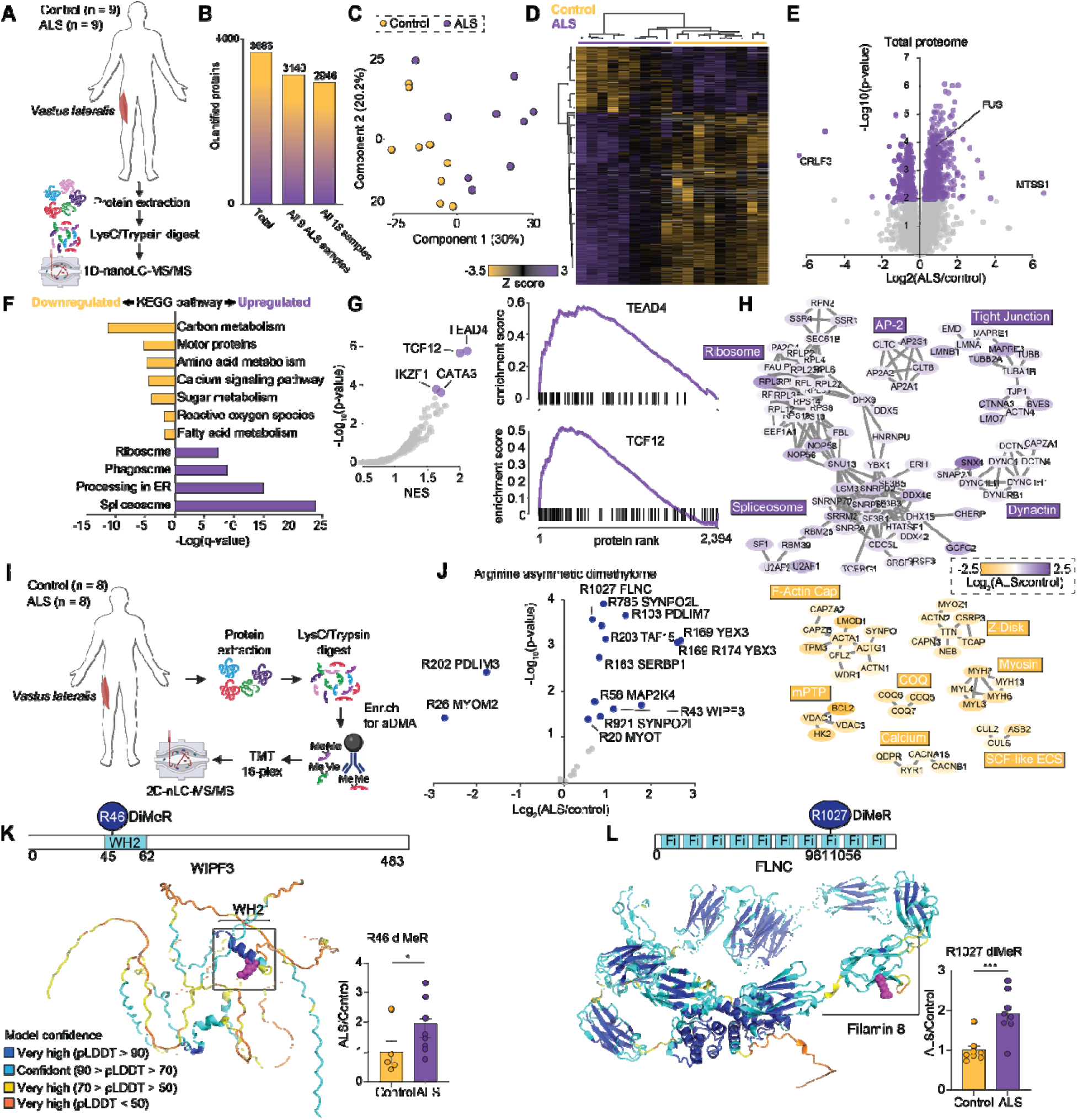
Quantification of the skeletal muscle proteome and asymmetric dimethyl proteome in human ALS. **A)** Overview of the experimental design to quantify the skeletal muscle proteome in ALS. **B)** Number of quantified proteins. **C)** Principal component analysis. **D)** Unsupervised hierarchal clustering. **E)** Proteome volcano plot. **F)** Kyoto encyclopaedia gene and genomes (KEGG) enrichment analysis. **(G)** CHEA3 geneset enrichment analysis. **(H)** Protein:protein interaction analysis from the STRINGdb. **I)** Overview of the experimental design to quantify the dimethyl proteome in ALS. **J)**. Dimethyl proteome volcano plot. Modification sites mapped onto alpha-fold predicted protein structures for WIPF3 **(K)** and FLNC **(L)**.

To quantify the skeletal muscle asymmetric dimethylome, LysC/tryptic peptides from the muscles of 8 patients with ALS and 8 control participants were enriched by immunoprecipitation using anti-aDMA antibodies, labeled with 16-plex tandem mass tags (TMT) and analyzed by 2D-nanoLC-MS/MS (**Figure 4I**). Subsequently, we normalized measured arginine methylated peptide ratios by the corresponding changes in protein abundance as determined above to investigate site-specific changes in methylation. We identified 43 unique asymmetric arginine methylation sites in 34 proteins, with 15 sites showing significant regulation of methylation state independent of protein abundance (**Figure 4J**) (**Table S3**). A further 8 dimethylated peptides were significantly increased in abundance, but we were unable to quantify their corresponding protein abundance (**Table S3**). The majority of the dimethylated peptides showed increased abundance in the ALS participants, consistent with the generalized global increase observed by western blot. Utilizing the PhosphoPlus database ^48^, none of the 15 sites have previously been assigned function, and among them, 5 sites were previously not identified using high or low throughput screening. The most upregulated site was R169 dimethylation on Y-box-binding protein 3 (YBX3), a repressor of immune cell function and maturation ^49^. While R26 dimethylation of MYOM2, an M-band structural protein responsible for sarcomere stability was the most downregulated ^50^. Moreover, we observed significant upregulation of dimethylation of RNA binding protein – TAF15 (TATA-box binding protein associated factor 15). Mutations in this protein have been previously associated with ALS pathogenesis ^51^. We also investigated the position of regulated arginine dimethylation sites within annotated protein domains mapped onto AlphaFold predicted structures and observed two other notable modification sites. Firstly, R46 dimethylation of WIPF3 (WAS/WASL-interacting protein family member 3), a cytoskeletal protein involved in actin binding, is positioned in its WH2 (Wiskott Aldrich syndrome homology region 2) motif, which participates in actin-subunit recruitment ^52^ (**Figure 4K**). The second modification is R1027 dimethylation of FLNC (filamin-C), a muscle-specific filamin that functions as a large actin-cross linker, is positioned in the 8^th^ (of 24) IgG-like repeat of a filamin (ABP280) rod domain, important for maintaining the structural integrity and function of the sarcomere ^53^ (**Figure 4L**). These data suggest that arginine dimethylation within known protein domains may regulate protein:protein interactions under this pathological setting of skeletal muscle atrophy.

Here, we present the largest compendium of protein changes in the skeletal muscle of human ALS participants and the first site-specific quantification of arginine dimethylation, which will be a valuable resource for further mechanistic studies.

### PRMT1 and aDMA overexpression provide resistance to fatigue and promote functional recovery

To further understand the functional consequences of asymmetric dimethylation in skeletal muscles, we selectively overexpressed PRMT1, the primary methyltransferase that deposits over 90% of the aDMA mark ^54^, in mice. We injected PRMT1 myoAAV into the TA and EDL muscles of 7 weeks old C57BL/6J mice (n = 10) in a paired experimental design, where the right leg was injected with PRMT1 myoAAV, while the left leg was injected with eGFP myoAAV as a control (**Figure 5A**). Four weeks after transduction, we observed significant increases in PRMT1 and aDMA levels in the right hindlimb muscles compared to the left (**Figure 5B-D**). There were no changes in muscle mass (**Figure 5E-F**) or cross-sectional area (CSA) (**Figure 5G**). We employed electrically induced contractions to EDL muscles *ex vivo*, revealing no changes in specific force (force generated normalized to CSA) (**Figure 5H**) or absolute tetanic force production (**Figure 5I**). Right EDL muscles expressing PRMT1 were more resistant to fatigue and exhibited greater functional recovery compared to left control EDL muscles (**Figure 5J**).

**Figure 5.**
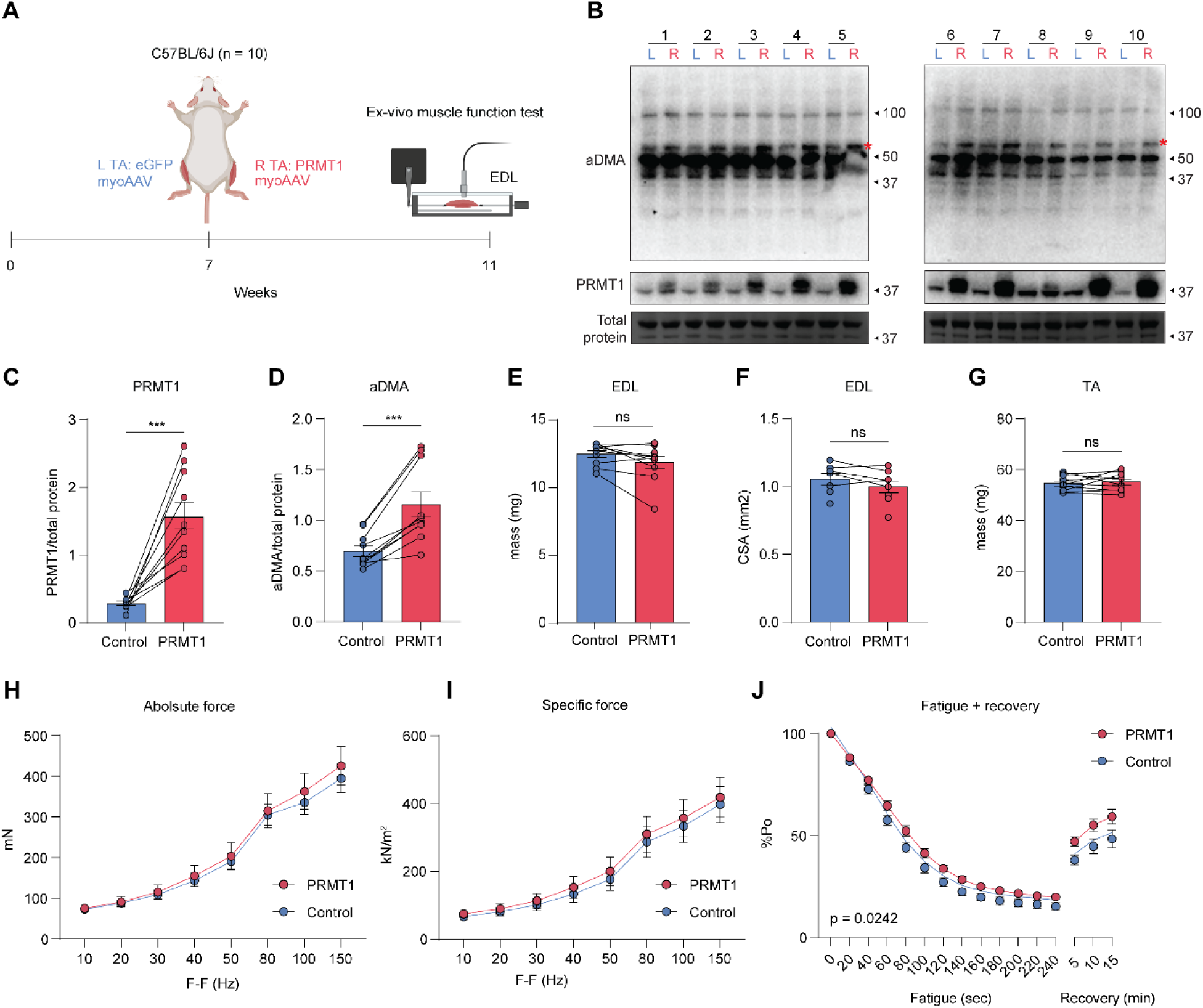
PRMT1 overexpression increases fatigue resistance and functional recovery. **A)**. Overview of the experimental design. **B-D)**. Western blot analysis (**B-C**) and quantification of PRMT1 (**C**) and aDMA (**D**) levels in left eGFP myoAAV control and right PRMT1 myoAAV TA muscles of 11 weeks old mice (n = 10). **E-G)**. Muscle mass of TA (**E**), EDL (**F**), and EDL cross-sectional area (CSA) (**G**) eGFP myoAAV control and PRMT1 myoAAV of mice (n = 10). **H-J)**. Specific force (**H)** and absolute tetanic force (**I**) production (mN) across incrementally increasing frequencies (10Hz – 150Hz, 350ms duration) in eGFP myoAAV control and PRMT1 myoAAV EDL muscles (n = 10). **J)**. Measurements of muscle fatigue and recovery by maximally stimulating the muscle once every 4 seconds for 4 minutes, followed by stimulation 5, 10, and 15 minutes after fatigue (n = 10). All quantified protein levels were normalized to total protein. Data are represented as mean ± SEM; ns = non-significant; α = 0.05; (B-E) Red asterisk denotes the protein band in which densitometry was performed (C-G) Paired t-test; (H-J) Two-way ANOVA.

## DISCUSSION

The regulation of arginine methylation under settings of muscle health and disease remains largely undefined. We investigated the regulation of arginine methylation in various models of health and disease, revealing a remodeling of asymmetric dimethylation (aDMA) of specific proteins in atrophy characterized by the loss of innervation, including muscle biopsies from human Amyotrophic lateral sclerosis (ALS) participants. To identify and quantify these modified proteins, we performed a comprehensive, integrated proteomic and arginine dimethylproteomic analysis of human ALS. Firstly, the proteomic results identified hundreds of novel changes and highlighted specific co-regulated protein complexes and pathways, providing a valuable resource to the field. Secondly, our study is the first to characterize the arginine dimethylome in human skeletal muscle, revealing novel site-specific changes in participants with ALS. We show that the overexpression of PRMT1 on aDMA resulted in increased fatigue resistance and functional recovery in mice, making this a potential therapeutic target in neurogenic atrophy.

In models of muscle health, arginine methylation showed subtle changes in acute human exercise (<2 fold) compared to the striking changes in protein phosphorylation (>10 fold), which we previously demonstrated in the exact same biopsies ^29^. Other works have similar reported limited changes in methylarginine/PRMTs following endurance/sprint exercise in humans and mice ^55,56^, overall suggesting a limited role of arginine methylation in acute exercise. In contrast, we observed significant increases in monomethyl-arginine (MMA)/aDMA in chronic muscle hypertrophy in mice. This differs from previous work, which reported subtle changes in arginine methylation following chronic spring training in humans ^56^. These differences are likely due to our use of a synergistic ablation (SA) mice model, which induces significantly greater muscular hypertrophy not typically seen in chronic exercise-induced adaptations ^57,58^. Taken together, the regulation of arginine methylation under chronic exercise settings and its role in physiological adaptations need to be further investigated.

Previous studies have demonstrated the potential role of arginine methylation in skeletal muscle metabolic diseases using a combination of cell culture and *in vivo* KO approaches ^16,17^. Here, we report limited changes in any of the three methylarginine variants in both insulin resistant and type 2 diabetes (T2D) models. We also observed no differences in the regulation of arginine methylation following acute stimulation of insulin for 30 min. Taken together, this suggests that although genetic manipulation of arginine methylation may modulate skeletal muscle insulin sensitivity, the levels of skeletal muscle arginine methylation do not appear to correlate with insulin-stimulated Akt signaling, insulin-stimulated glucose uptake or hyperglycemia.

We then investigated arginine methylation in skeletal muscle atrophy, revealing the regulation of aDMA in all settings characterized by the loss of innervation, including human ALS skeletal muscle biopsies. These changes occur most consistently at ∼60kDa, suggesting the existence of functionally conserved protein(s) important in neurogenic atrophy (see below). We observed an upregulation of aDMA in both skeletal muscle atrophy and hypertrophy, suggesting that aDMA functions as a protective mechanism for muscles, potentially via the upregulation of protein synthesis to combat atrophy-induced muscle loss and assist in hypertrophy-induced muscle gain. Indeed, it has recently been shown that PRMT1 and arginine methylation of NPRL2 are required for methionine/SAM-dependent activation of mTORC1 ^59^. These findings agree with previous studies where, generally, the knockdown/knockout of PRMTs resulted in decreases in muscle mass/function ^13,14^. In the future, the regulation of arginine methylation in additional atrophy settings such as sarcopenia and/or cancer cachexia should be explored.

We performed a comprehensive analysis of the skeletal muscle proteome/methyl proteome in participants with ALS. We report the largest compendium of protein changes in the skeletal muscle of human ALS with the identification of 793 differentially regulated proteins (**Table S1**). We present a dataset much greater than previous proteomic studies of ALS participants biopsies taken from skeletal muscle (e.g. 11 ^60^ and 20 ^61^ differentially regulated proteins) or other tissues, such as the spinal cord (281 ^62^ differentially proteins), representing a new resource to understand the pathological changes of ALS. With emerging evidence suggesting that the skeletal muscle can play an important role in ALS etiopathogenesis ^63^, our study provides a valuable proteomic resource as a mean to understand the role the skeletal muscle proteome plays in ALS. Additionally, we report the first comprehensive mass spectrometry-based annotation of the human asymmetric dimethyl proteome in ALS, including a subset of 43 unique dimethylation sites. Previous work has similarly quantified 117 aDMA sites in mice quadricep muscles ^64^. We initially expected to quantify a much larger set of methylation sites given that others have reported upwards of 1000 – 8000 MMA sites in HEK 293 cells ^18,65^. These differences could be due to the type of methylarginine enriched (i.e. MMA vs aDMA), the type of tissue investigated ^66^, and cell culture vs tissue differences (i.e. HEK 293 vs skeletal muscle) ^18,65,67^. Overall, our data revealed that all regulated methyl-proteins, except for one, were upregulated (consistent with the western blot analyses above), including WIPF3 (WAS/WASL Interacting Protein Family Member 3) and FLNC (Filamin C). Considering how a major consequence of muscle atrophy such as those seen in ALS is the degradation of sarcomeric and cytoskeleton proteins, we hypothesize that the upregulation of R46 WIPF3 may increase actin-subunit recruitment of the cytoskeleton, and R1027 FLNC may increase the structural integrity of the sarcomere, collectively protecting muscles against degradation by increasing the contractile frequency and stability of myofibers. As mentioned, we observed a common protein of ∼60Kda across all investigated atrophy models. To identify this protein, we matched our dimethyl proteome data within the mass range of ∼ 55-65Kda, revealing two regulated candidate proteins: R20 on MYOT (55.4Kda) and R203 on TAF15 (61.8Kda). We then interrogated a deep proteomic analysis of mouse skeletal muscle that included intensity-based absolute quantification (iBAQ) to rank their protein abundances ^68^. This allowed us to infer which protein is more abundant and, hence, more likely to be the prominent protein displayed in the western blot. Based on this analysis, this most abundant protein of approximately 60kDa, is MYOT (Myotilin) - a sarcomeric protein, which is ranked as the 83rd most abundant protein in the mouse skeletal muscle proteome. Future experiments are needed to better characterize this protein and the functional implications of arginine dimethylation on skeletal muscle atrophy. However, mutations in *MYOT* have previously been associated with myopathy ^69^.

Finally, we show that overexpression of PRMT1 – the primary enzyme responsible for over 90% of aDMA synthesis ^54^, resulted in increased fatigue resistance and functional recovery in adult mice skeletal muscles without any changes in muscle mass or function. We initially hypothesized that the overexpression of PRMT1 and aDMA might lead to an increase in muscle mass and contractile force, as muscle specific PRMT1 KO generated previously by Kang and colleagues showed a decrease in both parameters ^13^. These differences are likely due to our use of an AAV-overexpression instead of the Cre-LoxP-knockout model. Our model shows distinct advantages as it exclusively targets processes in adult skeletal muscle maintenance (i.e. protein turnover and muscle regeneration) without affecting processes involved in muscle development (i.e. myoblast formation, myotube formation) ^70–72^. This is important as the use of therapeutics in muscles to increase mass/function will need to be given post-birth and not during embryogenesis. Our results provide a rationale for pursuing PRMT1 as a therapeutic target in diseases such as ALS and muscular dystrophies where muscle fatigue is common ^73–75^. Future experiments are needed to explore if the same protective effect can be achieved in models of ALS.

In summary, we investigated the regulation of arginine methylation in various models of skeletal muscle health and disease, revealing an upregulation of asymmetric dimethylation in models of atrophy characterized by the loss of innervation, including in patients with ALS. We performed an integrated proteomics/methyl proteomics analysis to quantify site-specific changes in aDMA in skeletal biopsies from human participants with ALS. We provide the first comprehensive mass spectrometry-based annotation of the human skeletal muscle proteome and dimethyl proteome in ALS, including the identification of 793 regulated proteins, 43 unique dimethylation sites, and novel site-specific changes in key proteins of the sarcomere and cytoskeleton. Finally, we show that overexpression of PRMT1 on aDMA resulted in increased fatigue resistance and functional recovery in mice. Our study provides evidence for asymmetric dimethylation as a regulator of muscle function and presents a valuable proteomics resource and rationale for numerous methylated and non-methylated proteins, including PRMT1, to be pursued for therapeutic development in ALS.

## DATA AVAILABILITY

The mass spectrometry proteomics data have been deposited to the ProteomeXchange Consortium via the PRIDE [1] partner repository with the dataset identifier PXD047184 (Username; Password: NbjNdPzb)

## AUTHOR CONTRIBUTIONS

J.P.H.W., R.B., Y.-N.K., C.A.G., M.K.M, K.I.W., C.S.C., B.K., E.A.R., P.J.C., F.J.S., S.T.N., B.L.P. performed experiments. J.P.H.W. and B.L.P. analyzed the data and wrote the manuscript. M.J.W., B.K., P.J.C., F.J.S., S.T.N. and B.L.P. supervised the work.

## ACKNOWLEDGMENTS

We thank Nicholas Williamson, Ching-Seng Ang, Shuai Nie, Swati Varshney and Michael Leeming for instrument support in the Bio21 Mass Spectrometry and Proteomics Facility. We also thank Bente Kiens and staff August Krogh Section for Molecular Physiology, Department of Nutrition, Exercise and Sports, Faculty of Science, The University of Copenhagen, Denmark for assistance with the exercise studies. This research was supported by access to the Melbourne Mouse Metabolic Phenotyping Platform at The University of Melbourne. This work was funded by an NHMRC Emerging Leader Investigator Grant (APP2009642) and a University of Melbourne Driving Research Momentum Grant to B.L.P. This work was also supported by a Department of Anatomy and Physiology (The University of Melbourne) ECR Seeding Grant to R.B. Y.-K.N. is a recipient of a School of Biomedical Science Postgraduate Award.

## REFERENCES

1. Frontera WR, Ochala J. Skeletal muscle: a brief review of structure and function. Calcif Tissue Int. Mar 2015;96(3):183–95. doi:10.1007/s00223-014-9915-y

2. Jackman RW, Kandarian SC. The molecular basis of skeletal muscle atrophy. Am J Physiol Cell Physiol. Oct 2004;287(4):C834–43. doi:10.1152/ajpcell.00579.2003

3. Khan MAB, Hashim MJ, King JK, Govender RD, Mustafa H, Al Kaabi J. Epidemiology of Type 2 Diabetes - Global Burden of Disease and Forecasted Trends. J Epidemiol Glob Health. Mar 2020;10(1):107–111. doi:10.2991/jegh.k.191028.001

4. Bostock EL, O’Dowd DN, Payton CJ, et al. The Effects of Resistance Exercise Training on Strength and Functional Tasks in Adults With Limb-Girdle, Becker, and Facioscapulohumeral Dystrophies. Front Neurol. 2019;10:1216. doi:10.3389/fneur.2019.01216

5. Little JP, Phillips SM. Resistance exercise and nutrition to counteract muscle wasting. Appl Physiol Nutr Metab. Oct 2009;34(5):817–28. doi:10.1139/h09-093

6. Laurin JL, Reid JJ, Lawrence MM, Miller BF. Long-term aerobic exercise preserves muscle mass and function with age. Current Opinion in Physiology. 2019/08/01/ 2019;10:70–74. 10.1016/j.cophys.2019.04.019

7. Chen YL, Ma YC, Tang J, et al. Physical exercise attenuates age-related muscle atrophy and exhibits anti-ageing effects via the adiponectin receptor 1 signalling. J Cachexia Sarcopenia Muscle. Aug 2023;14(4):1789–1801. doi:10.1002/jcsm.13257

8. Richter EA, Mikines KJ, Galbo H, Kiens B. Effect of exercise on insulin action in human skeletal muscle. J Appl Physiol (1985). Feb 1989;66(2):876–85. doi:10.1152/jappl.1989.66.2.876

9. Nystoriak MA, Bhatnagar A. Cardiovascular Effects and Benefits of Exercise. Front Cardiovasc Med. 2018;5:135. doi:10.3389/fcvm.2018.00135

10. Booth FW, Roberts CK, Laye MJ. Lack of exercise is a major cause of chronic diseases. Compr Physiol. Apr 2012;2(2):1143–211. doi:10.1002/cphy.c110025

11. Blanc RS, Richard S. Arginine Methylation: The Coming of Age. Mol Cell. Jan 5 2017;65(1):8–24. doi:10.1016/j.molcel.2016.11.003

12. vanLieshout TL, Ljubicic V. The emergence of protein arginine methyltransferases in skeletal muscle and metabolic disease. Am J Physiol Endocrinol Metab. Dec 1 2019;317(6):E1070–e1080. doi:10.1152/ajpendo.00251.2019

13. Choi S, Jeong HJ, Kim H, et al. Skeletal muscle-specific Prmt1 deletion causes muscle atrophy via deregulation of the PRMT6-FOXO3 axis. Autophagy. Jun 2019;15(6):1069–1081. doi:10.1080/15548627.2019.1569931

14. Stouth DW, vanLieshout TL, Ng SY, et al. CARM1 Regulates AMPK Signaling in Skeletal Muscle. iScience. Nov 20 2020;23(11):101755. doi:10.1016/j.isci.2020.101755

15. Zhang T, Günther S, Looso M, et al. Prmt5 is a regulator of muscle stem cell expansion in adult mice. Nature Communications. 2015/06/01 2015;6(1):7140. doi:10.1038/ncomms8140

16. Jeong HJ, Lee HJ, Vuong TA, et al. Prmt7 Deficiency Causes Reduced Skeletal Muscle Oxidative Metabolism and Age-Related Obesity. Diabetes. Jul 2016;65(7):1868–82. doi:10.2337/db15-1500

17. Iwasaki H, Yada T. Protein arginine methylation regulates insulin signaling in L6 skeletal muscle cells. Biochem Biophys Res Commun. Dec 28 2007;364(4):1015–21. doi:10.1016/j.bbrc.2007.10.113

18. Larsen SC, Sylvestersen KB, Mund A, et al. Proteome-wide analysis of arginine monomethylation reveals widespread occurrence in human cells. Sci Signal. Aug 30 2016;9(443):rs9. doi:10.1126/scisignal.aaf7329

19. vanLieshout TL, Stouth DW, Hartel NG, et al. The CARM1 transcriptome and arginine methylproteome mediate skeletal muscle integrative biology. Molecular Metabolism. 2022/10/01/ 2022;64:101555. 10.1016/j.molmet.2022.101555

20. Kjøbsted R, Kido K, Larsen JK, et al. Measurement of Insulin- and Contraction-Stimulated Glucose Uptake in Isolated and Incubated Mature Skeletal Muscle from Mice. J Vis Exp. May 16 2021;(171)doi:10.3791/61398

21. Tabbaa M, Gomez TR, Campelj DG, Gregorevic P, Hayes A, Goodman CA. The regulation of polyamine pathway proteins in models of skeletal muscle hypertrophy and atrophy: a potential role for mTORC1. American Journal of Physiology-Cell Physiology. 2021;320(6):C987–C999. doi:10.1152/ajpcell.00078.2021

22. Wong PC, Pardo CA, Borchelt DR, et al. An adverse property of a familial ALS-linked SOD1 mutation causes motor neuron disease characterized by vacuolar degeneration of mitochondria. Neuron. Jun 1995;14(6):1105–16. doi:10.1016/0896-6273(95)90259-7

23. Arnold ES, Ling SC, Huelga SC, et al. ALS-linked TDP-43 mutations produce aberrant RNA splicing and adult-onset motor neuron disease without aggregation or loss of nuclear TDP-43. Proc Natl Acad Sci U S A. Feb 19 2013;110(8):E736–45. doi:10.1073/pnas.1222809110

24. Roberts BR, Lim NK, McAllum EJ, et al. Oral treatment with Cu(II)(atsm) increases mutant SOD1 in vivo but protects motor neurons and improves the phenotype of a transgenic mouse model of amyotrophic lateral sclerosis. J Neurosci. Jun 4 2014;34(23):8021–31. doi:10.1523/jneurosci.4196-13.2014

25. Brooks BR, Miller RG, Swash M, Munsat TL. El Escorial revisited: revised criteria for the diagnosis of amyotrophic lateral sclerosis. Amyotroph Lateral Scler Other Motor Neuron Disord. Dec 2000;1(5):293–9. doi:10.1080/146608200300079536

26. Ioannides ZA, Steyn FJ, Henderson RD, McCombe PA, Ngo ST. Anthropometric measures are not accurate predictors of fat mass in ALS. Amyotroph Lateral Scler Frontotemporal Degener. Nov 2017;18(7-8):486–491. doi:10.1080/21678421.2017.1317811

27. Steyn FJ, Ioannides ZA, van Eijk RPA, et al. Hypermetabolism in ALS is associated with greater functional decline and shorter survival. J Neurol Neurosurg Psychiatry. Oct 2018;89(10):1016–1023. doi:10.1136/jnnp-2017-317887

28. Tarnopolsky MA, Pearce E, Smith K, Lach B. Suction-modified Bergström muscle biopsy technique: experience with 13,500 procedures. Muscle Nerve. May 2011;43(5):717–25. doi:10.1002/mus.21945

29. Blazev R, Carl CS, Ng YK, et al. Phosphoproteomics of three exercise modalities identifies canonical signaling and C18ORF25 as an AMPK substrate regulating skeletal muscle function. Cell Metab. Oct 4 2022;34(10):1561–1577.e9. doi:10.1016/j.cmet.2022.07.003

30. Le Couteur DG, Solon-Biet SM, Parker BL, et al. Nutritional reprogramming of mouse liver proteome is dampened by metformin, resveratrol, and rapamycin. Cell Metab. Dec 7 2021;33(12):2367–2379.e4. doi:10.1016/j.cmet.2021.10.016

31. Bruderer R, Bernhardt OM, Gandhi T, et al. Optimization of Experimental Parameters in Data-Independent Mass Spectrometry Significantly Increases Depth and Reproducibility of Results. Mol Cell Proteomics. Dec 2017;16(12):2296–2309. doi:10.1074/mcp.RA117.000314

32. Käll L, Canterbury JD, Weston J, Noble WS, MacCoss MJ. Semi-supervised learning for peptide identification from shotgun proteomics datasets. Nat Methods. Nov 2007;4(11):923–5. doi:10.1038/nmeth1113

33. Taus T, Köcher T, Pichler P, et al. Universal and confident phosphorylation site localization using phosphoRS. J Proteome Res. Dec 2 2011;10(12):5354–62. doi:10.1021/pr200611n

34. Tyanova S, Temu T, Sinitcyn P, et al. The Perseus computational platform for comprehensive analysis of (prote)omics data. Nat Methods. Sep 2016;13(9):731–40. doi:10.1038/nmeth.3901

35. Zhou Y, Zhou B, Pache L, et al. Metascape provides a biologist-oriented resource for the analysis of systems-level datasets. Nat Commun. Apr 3 2019;10(1):1523. doi:10.1038/s41467-019-09234-6

36. Keenan AB, Torre D, Lachmann A, et al. ChEA3: transcription factor enrichment analysis by orthogonal omics integration. Nucleic Acids Res. Jul 2 2019;47(W1):W212–w224. doi:10.1093/nar/gkz446

37. Molendijk J, Yip R, Parker BL. urPTMdb/TeaProt: Upstream and Downstream Proteomics Analysis. J Proteome Res. Feb 3 2023;22(2):302–310. doi:10.1021/acs.jproteome.2c00048

38. Schneider CA, Rasband WS, Eliceiri KW. NIH Image to ImageJ: 25 years of image analysis. Nature Methods. 2012/07/01 2012;9(7):671–675. doi:10.1038/nmeth.2089

39. Tabebordbar M, Lagerborg KA, Stanton A, et al. Directed evolution of a family of AAV capsid variants enabling potent muscle-directed gene delivery across species. Cell. Sep 16 2021;184(19):4919–4938.e22. doi:10.1016/j.cell.2021.08.028

40. Khan IF, Hirata RK, Russell DW. AAV-mediated gene targeting methods for human cells. Nat Protoc. Apr 2011;6(4):482–501. doi:10.1038/nprot.2011.301

41. Goodman CA, Mabrey DM, Frey JW, et al. Novel insights into the regulation of skeletal muscle protein synthesis as revealed by a new nonradioactive in vivo technique. Faseb j. Mar 2011;25(3):1028–39. doi:10.1096/fj.10-168799

42. Coleman DL. Obese and diabetes: two mutant genes causing diabetes-obesity syndromes in mice. Diabetologia. Mar 1978;14(3):141–8. doi:10.1007/bf00429772

43. Watt KI, Turner BJ, Hagg A, et al. The Hippo pathway effector YAP is a critical regulator of skeletal muscle fibre size. Nature Communications. 2015/01/12 2015;6(1):6048. doi:10.1038/ncomms7048

44. Blokhuis AM, Groen EJ, Koppers M, van den Berg LH, Pasterkamp RJ. Protein aggregation in amyotrophic lateral sclerosis. Acta Neuropathol. Jun 2013;125(6):777–94. doi:10.1007/s00401-013-1125-6

45. Logroscino G, Piccininni M. Amyotrophic Lateral Sclerosis Descriptive Epidemiology: The Origin of Geographic Difference. Neuroepidemiology. 2019;52(1-2):93–103. doi:10.1159/000493386

46. Kanehisa M, Goto S. KEGG: kyoto encyclopedia of genes and genomes. Nucleic Acids Res. Jan 1 2000;28(1):27–30. doi:10.1093/nar/28.1.27

47. Szklarczyk D, Franceschini A, Wyder S, et al. STRING v10: protein-protein interaction networks, integrated over the tree of life. Nucleic Acids Res. Jan 2015;43(Database issue):D447–52. doi:10.1093/nar/gku1003

48. Hornbeck PV, Zhang B, Murray B, Kornhauser JM, Latham V, Skrzypek E. PhosphoSitePlus, 2014: mutations, PTMs and recalibrations. Nucleic Acids Res. Jan 2015;43(Database issue):D512–20. doi:10.1093/nar/gku1267

49. Shi Y, Liu CH, Roberts AI, et al. Granulocyte-macrophage colony-stimulating factor (GM-CSF) and T-cell responses: what we do and don’t know. Cell Res. Feb 2006;16(2):126–33. doi:10.1038/sj.cr.7310017

50. Lamber EP, Guicheney P, Pinotsis N. The role of the M-band myomesin proteins in muscle integrity and cardiac disease. Journal of Biomedical Science. 2022/03/07 2022;29(1):18. doi:10.1186/s12929-022-00801-6

51. Kapeli K, Pratt GA, Vu AQ, et al. Distinct and shared functions of ALS-associated proteins TDP-43, FUS and TAF15 revealed by multisystem analyses. Nat Commun. Jul 5 2016;7:12143. doi:10.1038/ncomms12143

52. Dominguez R. The WH2 Domain and Actin Nucleation: Necessary but Insufficient. Trends Biochem Sci. Jun 2016;41(6):478–490. doi:10.1016/j.tibs.2016.03.004

53. Mao Z, Nakamura F. Structure and Function of Filamin C in the Muscle Z-Disc. Int J Mol Sci. Apr 13 2020;21(8)doi:10.3390/ijms21082696

54. Tang J, Kao PN, Herschman HR. Protein-arginine methyltransferase I, the predominant protein-arginine methyltransferase in cells, interacts with and is regulated by interleukin enhancer-binding factor 3. J Biol Chem. Jun 30 2000;275(26):19866–76. doi:10.1074/jbc.M000023200

55. Vanlieshout TL, Stouth DW, Tajik T, Ljubicic V. Exercise-induced Protein Arginine Methyltransferase Expression in Skeletal Muscle. Med Sci Sports Exerc. Mar 2018;50(3):447–457. doi:10.1249/mss.0000000000001476

56. vanLieshout TL, Bonafiglia JT, Gurd BJ, Ljubicic V. Protein arginine methyltransferase biology in humans during acute and chronic skeletal muscle plasticity. Journal of Applied Physiology. 2019;127(3):867–880. doi:10.1152/japplphysiol.00142.2019

57. Terena SM, Fernandes KP, Bussadori SK, Deana AM, Mesquita-Ferrari RA. Systematic review of the synergist muscle ablation model for compensatory hypertrophy. Rev Assoc Med Bras (1992). Feb 2017;63(2):164–172. doi:10.1590/1806-9282.63.02.164

58. Hubbard RW, Ianuzzo CD, Mathew WT, Linduska JD. Compensatory adaptations of skeletal muscle composition to a long-term functional overload. Growth. Mar 1975;39(1):85–93.

59. Jiang C, Liu J, He S, et al. PRMT1 orchestrates with SAMTOR to govern mTORC1 methionine sensing via Arg-methylation of NPRL2. Cell Metab. Dec 5 2023;35(12):2183–2199.e7. doi:10.1016/j.cmet.2023.11.001

60. Elf K, Shevchenko G, Nygren I, et al. Alterations in muscle proteome of patients diagnosed with amyotrophic lateral sclerosis. J Proteomics. Aug 28 2014;108:55–64. doi:10.1016/j.jprot.2014.05.004

61. Conti A, Riva N, Pesca M, et al. Increased expression of Myosin binding protein H in the skeletal muscle of amyotrophic lateral sclerosis patients. Biochim Biophys Acta. Jan 2014;1842(1):99–106. doi:10.1016/j.bbadis.2013.10.013

62. Iridoy MO, Zubiri I, Zelaya MV, et al. Neuroanatomical Quantitative Proteomics Reveals Common Pathogenic Biological Routes between Amyotrophic Lateral Sclerosis (ALS) and Frontotemporal Dementia (FTD). Int J Mol Sci. Dec 20 2018;20(1)doi:10.3390/ijms20010004

63. Shefner JM, Musaro A, Ngo ST, et al. Skeletal muscle in amyotrophic lateral sclerosis. Brain. 2023;146(11):4425–4436. doi:10.1093/brain/awad202

64. vanLieshout TL, Stouth DW, Hartel NG, et al. The CARM1 transcriptome and arginine methylproteome mediate skeletal muscle integrative biology. Mol Metab. Oct 2022;64:101555. doi:10.1016/j.molmet.2022.101555

65. Li W-j, He Y-h, Yang J-j, et al. Profiling PRMT methylome reveals roles of hnRNPA1 arginine methylation in RNA splicing and cell growth. Nature Communications. 2021/03/29 2021;12(1):1946. doi:10.1038/s41467-021-21963-1

66. Onwuli DO, Rigau-Roca L, Cawthorne C, Beltran-Alvarez P. Mapping arginine methylation in the human body and cardiac disease. PROTEOMICS – Clinical Applications. 2017;11(1-2):1600106. 10.1002/prca.201600106

67. Lim Y, Lee JY, Ha SJ, Yu S, Shin JK, Kim HC. Proteome-wide identification of arginine methylation in colorectal cancer tissues from patients. Proteome Science. 2020/05/19 2020;18(1):6. doi:10.1186/s12953-020-00162-8

68. Deshmukh AS, Murgia M, Nagaraj N, Treebak JT, Cox J, Mann M. Deep proteomics of mouse skeletal muscle enables quantitation of protein isoforms, metabolic pathways, and transcription factors. Mol Cell Proteomics. Apr 2015;14(4):841–53. doi:10.1074/mcp.M114.044222

69. Guglielmi V, Pancheri E, Cannone E, et al. A novel in-frame deletion in MYOT causes an early adult onset distal myopathy. Clin Genet. Dec 2023;104(6):705–710. doi:10.1111/cge.14413

70. Romero NB, Mezmezian M, Fidziańska A. Main steps of skeletal muscle development in the human: morphological analysis and ultrastructural characteristics of developing human muscle. Handb Clin Neurol. 2013;113:1299–310. doi:10.1016/b978-0-444-59565-2.00002-2

71. Bentzinger CF, Wang YX, Rudnicki MA. Building muscle: molecular regulation of myogenesis. Cold Spring Harb Perspect Biol. Feb 1 2012;4(2)doi:10.1101/cshperspect.a008342

72. Musarò A. Muscle Homeostasis and Regeneration: From Molecular Mechanisms to Therapeutic Opportunities. Cells. Sep 4 2020;9(9)doi:10.3390/cells9092033

73. El-Aloul B, Speechley KN, Wei Y, Wilk P, Campbell C. Fatigue in young people with Duchenne muscular dystrophy. Dev Med Child Neurol. Feb 2020;62(2):245–251. doi:10.1111/dmcn.14248

74. Angelini C, Tasca E. Fatigue in muscular dystrophies. Neuromuscul Disord. Dec 2012;22 Suppl 3(3-3):S214–20. doi:10.1016/j.nmd.2012.10.010

75. Gibbons CJ, Thornton EW, Young CA. The patient experience of fatigue in motor neurone disease. Front Psychol. 2013;4:788. doi:10.3389/fpsyg.2013.00788

